# The G protein-biased PZM21 and TRV130 act as partial agonists of μ-opioid receptors signaling to ion channel targets

**DOI:** 10.1101/445536

**Authors:** Yevgen Yudin, Tibor Rohacs

## Abstract

Opioids exert many of their acute effects through modulating ion channels via G_βγ_ subunits. Some of their side effects are attributed to β-arrestin recruitment, and several biased agonists that do not activate this pathway have been developed recently. Here we tested the effects of TRV130, PZM21 and herkinorin, three G-protein biased agonists of μ-opioid receptors (μOR), on ion channel targets. Compared to the full μOR agonist DAMGO, all three biased agonists induced smaller activation of G protein-coupled inwardly rectifying potassium channels (GIRK2), and smaller inhibition of Transient Receptor Potential Melastatin (TRPM3) channels. Furthermore, co-application of TRV130 or PZM21, but not herkinorin reduced the effects of DAMGO on both ion channels. Ca_V_2.2 was also inhibited less by PZM21 and TRV130 than by DAMGO. TRV130, PZM21 and herkinorin were also less effective than DAMGO in inducing dissociation of the G_αi_ /G_βγ_ complex. We conclude that TRV130, PZM21 are partial agonists of μOR.

## INTRODUCTION

Chronic pain is an unsolved medical problem, causing immense suffering to millions of people worldwide (Basbaum et al., 2009). Opioids remain the main therapy against severe chronic pain, but these medications cause multiple side effects limiting their application, and are also highly addictive. The lack of better pain control is believed to be a significant driving force behind the opioid addiction crisis (Skolnick and Volkow, 2016).

Opioids exert their effects via binding to μ-δ- or κ-opioid receptors (Stein, 2016). All these receptors act via inhibitory G-proteins in the G_αi/o_ family. Upon receptor activation, the G_α_ subunits dissociate from G_βγ_, and both subunits activate multiple downstream effectors. Opioid receptor activation also induces recruitment of β-arrestins, leading to receptor desensitization, internalization and various other effects. β-arrestins however, are not only important for desensitization, but they are also emerging as independent signaling mediators regulating various effectors (DeWire et al., 2007), including ion channels (Liu et al., 2017). Most clinically useful analgesic effects of opioids are mediated by μ-opioid receptors (μOR). Some side effects of opioids, such as tolerance have been attributed μOR signaling via β-arrestins (Raehal et al., 2011).

On the cellular level, opioid receptor activation leads to decreased cAMP production, an effect mediated by inhibition of the adenylate cyclase enzyme by G_αi_. Many acute effects of opioids are mediated by reduction of neuronal activity via modulation of ion channels by G_βγ_ subunits. The two classical actions of G_αi/o_ signaling are activation of G-protein coupled inwardly rectifying K^+^ channels (GIRK) and inhibition of N- and P/Q type voltage gated Ca^2+^ channels (Ca_v_2.2 and Ca_v_2.1) (Clapham and Neer, 1997), both effects leading to reduced neuronal activity.

We and others recently identified Transient Receptor Potential Melastatin 3 (TRPM3) as a novel ion channel target of G_βγ_ subunits; these channels are robustly inhibited by activation of a variety of G_αi_-coupled receptors, including μOR in dorsal root ganglion (DRG) neurons, as well as in expression systems (Badheka et al., 2017; Dembla et al., 2017; Quallo et al., 2017). TRPM3 is a heat-activated Ca^2+^ permeable non-selective cation channel; its genetic deletion in mice results in reduced heat sensitivity (Vriens et al., 2011; Vandewauw et al., 2018). TRPM3 also has chemical activators such as the endogenous neurosteroid Pregnenolone Sulphate (PregS) (Wagner et al., 2008) as well as synthetic chemical activators including CIM0216 and nifedipine (Held et al., 2015). This channel is also involved in regulating glucose homeostasis and expressed in many other tissues including pancreatic beta cells, retina and brain (Oberwinkler and Philipp, 2014).

Several biased agonists of μOR that activate G-protein signaling, but do not lead to β-arrestin recruitment have been recently described with the goal of developing painkillers with reduced side effects. Herkinorin was the first G-protein-biased agonist that was claimed not to lead to β-arrestin recruitment and receptor internalization (Groer et al., 2007). TRV130 (oliceridine) developed by Trevena pharmaceuticals (DeWire et al., 2007), has shown promising analgesic effects in clinical trials (Viscusi et al., 2016; Singla et al., 2017). The latest G-protein biased μOR agonist PZM21 was developed based on structure guided computational modeling, and showed promising analgesic results in mice (Manglik et al., 2016). The study showed that PZM21 as well as TRV130 did not lead to β-arrestin recruitment, but found that herkinorin evoked β-arrestin recruitment similar to that induced by full agonists (Manglik et al., 2016). The effects of these G-protein biased agonists have not yet been tested on ion channel targets of G_αi_ signaling.

Here we tested the effects of the G-protein biased μOR agonists TRV130, PZM21 and herkinorin on three ion channel targets of G_βγ_. Surprisingly we found that all three biased agonists induced significantly smaller inhibition of TRPM3, and Ca_V_2.2 voltage gated Ca^2+^ channels and smaller activation of GIRK2 than the full agonist DAMGO. Furthermore, PZM21 and TRV130, but not herkinorin relieved DAMGO-induced inhibition of TRPM3, and reduced DAMGO-induced activation of GIRK2 channels. All three G-protein biased agonists were less efficient in inducing G-protein dissociation than DAMGO. Overall our data indicate that PZM21 and TRV130 act as partial μOR agonists.

## RESULTS

### G-protein biased μOR agonists inhibit TRPM3 channels less than the full agonist DAMGO

The initial motivation of this study was to test the potential involvement of the β-arrestin pathway in the inhibition of TRPM3 channels. To this end, first we tested the effects of PZM21, TRV130 and Herkinorin on Ca^2+^ signals induced by the TRPM3 agonist PregS in DRG neurons. We have found that the full μOR agonist DAMGO (1 μM) induced strong inhibition in the vast majority of the PregS responsive neurons (61 of 64; 95%) (Fig. 1A), while application of the other three agonists at the same concentration affected only a much smaller fraction of the PregS responsive neurons (Fig 1B-C). For 1 μM Herkinorin we observed inhibition in 38 out of 69 (55 %) tested neurons (Fig. 1B), for 1 μM PZM21 in 14 out of 22 (63 %) (Fig. 1C), and for 1 μM TRV130 in 12 out of 22 neurons (54 %) (Fig. 1D). We also observed that the inhibitory effects of these three biased agonists were longer lasting compared to the quickly reversible effects of DAMGO.

**Fig 1.**
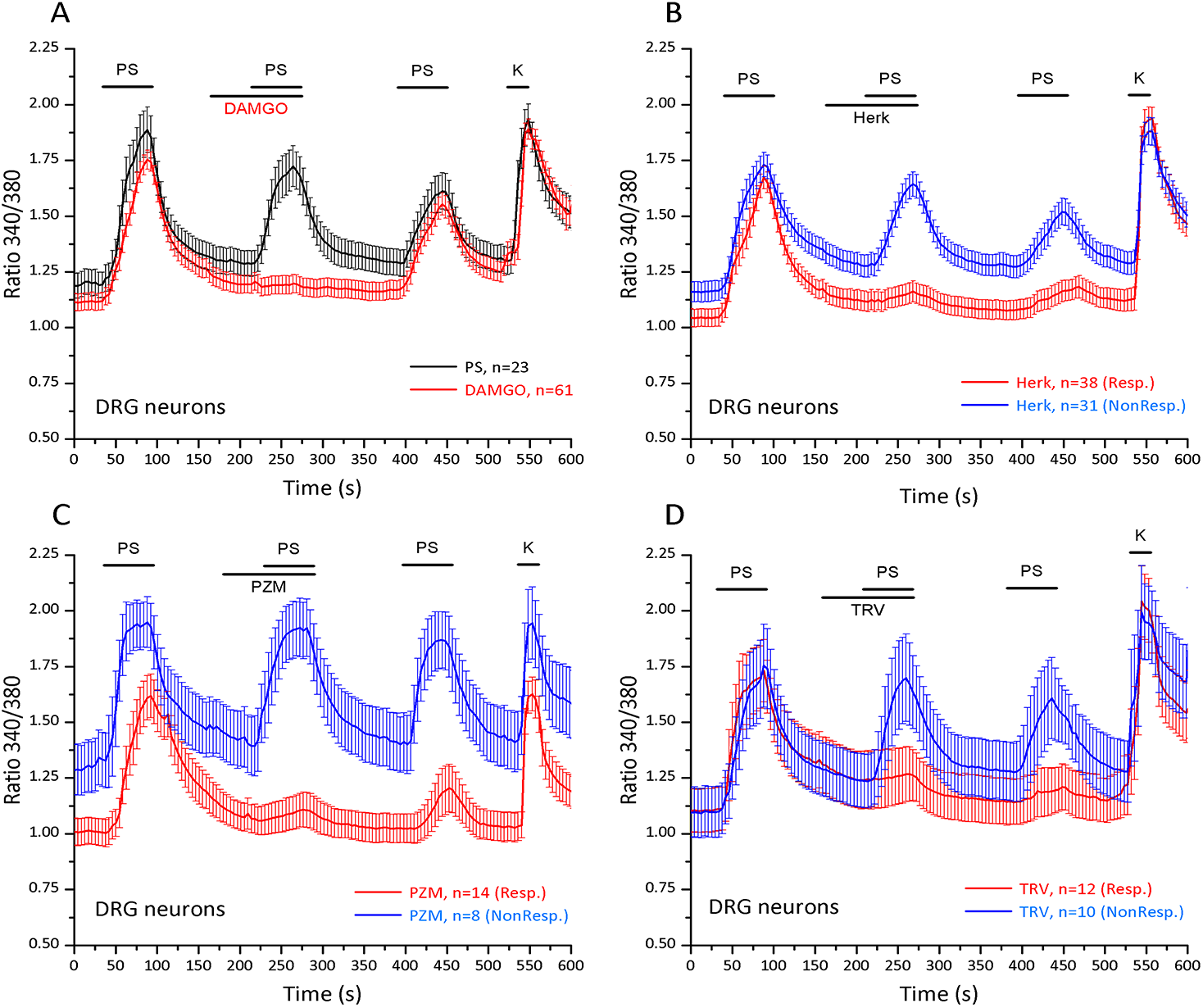
Differential inhibition of PregS-induced Ca^2^+ signals by μOR agonists in DRG neurons. Ca^2^+ imaging experiments in Fura-2 loaded mouse DRG neurons were performed as described in Materials and methods. **(A)** Black trace shows mean ± SEM of the effect of three consecutive applications of 12.5 μM PregS (PS) from neurons responsive to this compound, n=23; 30 mM KCl was applied at the end of the experiment. In red color 1 μM DAMGO was applied before the second application of PregS, out of 64 neurons inhibition was observed in 61, the non-responding cells were not included in the average trace. **(B)** Similar measurements with the application of 1 μM Herkinorin; the two traces show the average ratios ± SEM in cells in which PregS-induced Ca^2+^ signals were inhibited by Herkinorin (red, n=38) and in cells where inhibition was not observed (blue, n=31). **(C)** Similar measurements with 1 μM PZM21 ; the two traces show the average ratios ± SEM in cells where PregS-induced Ca^2+^ signal was inhibited by PZM21 (red, n=14) and in cells where it was not inhibited (blue, n=8). **(D)** Similar measurements with 1 μM TRV130; the two traces show the average ratios ± SEM in cells where PregS-induced Ca^2+^ signals were inhibited by TRV130 (red, n=12) and in cells where they were not inhibited (blue, n=10).

DRG neurons express all three types of opioid receptors − μ, δ, κ, which may potentially confound interpretation of these data, as some of the μOR agonists may cross-react with other opioid receptors. Therefore we co-expressed μOR and TRPM3 channels in HEK cells and performed Ca^2+^ imaging experiments similar to those in DRG neurons. We found that similar to DRG neurons, PZM1, TRV130 and herkinorin evoked significantly smaller inhibition of PregS-induced Ca^2+^ signals than DAMGO (Fig 2). DAMGO (0.5 μM) evoked, on average, an 89 % inhibition of the PregS-induced Ca^2+^ signal, for 1 μM PZM21 average inhibition was 29 %, for 1 μM TRV130 it was 20 %, and for 1 μM Herkinorin it was 37 %. Also the number of the cell where inhibition was more than 20 % of the PregS-induced Ca^2+^ signal was much higher for DAMGO (207 out of 208 cells, 99.5 %) than for Herkinorin (96 out of 173; 55.5 %) or for TRV130 (76 out of 227; 33.5 %) or for PZM21 (98 out of 187; 52.4 %).

**Fig 2.**
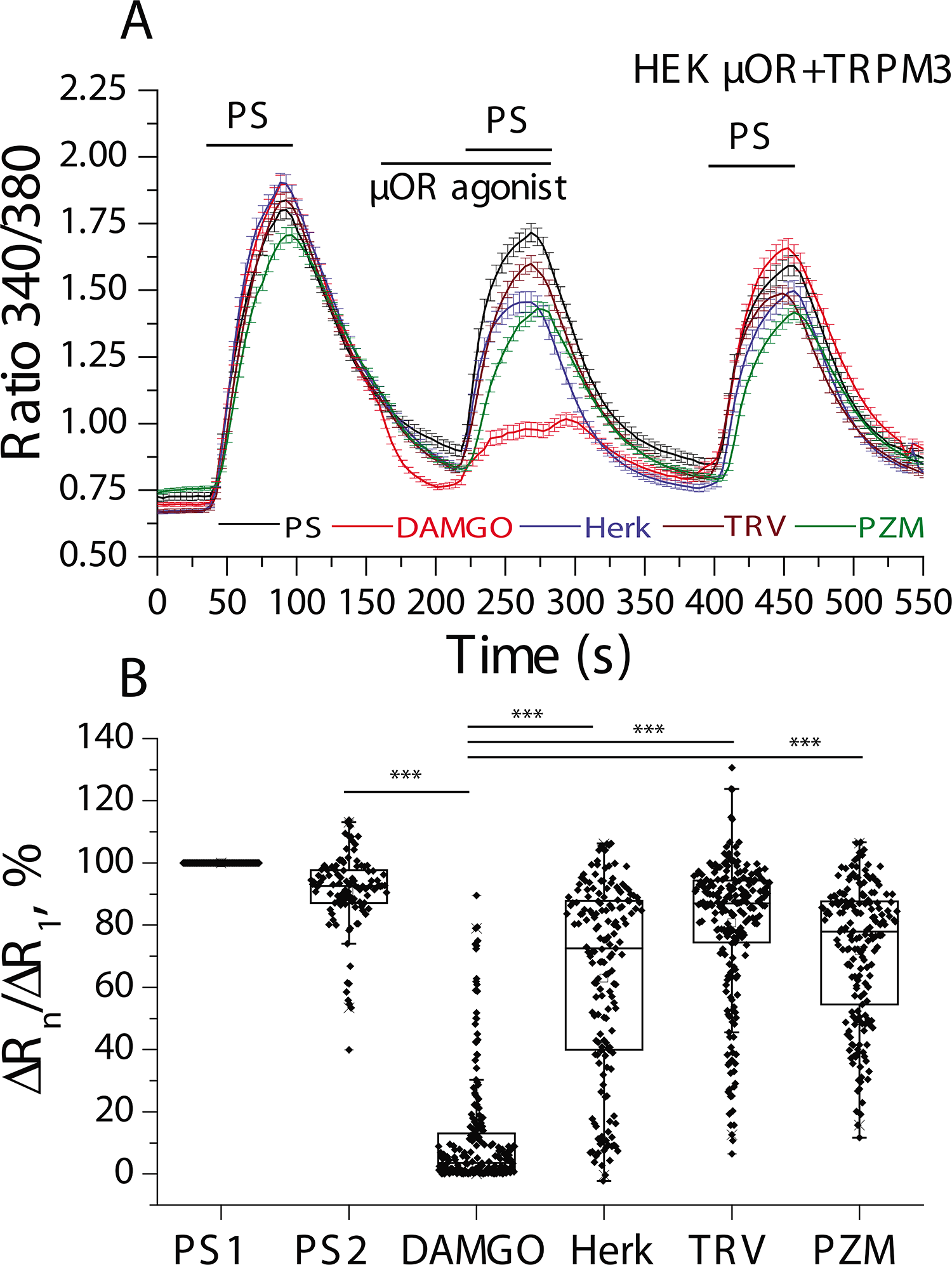
Differential inhibition of PregS-induced Ca^2^+ signals by μOR agonists in HEK293 cells. Ca^2+^ imaging experiments in HEK cells transfected with mTRPM3 and μOR were performed as described in Materials and methods. **(A)** Traces show mean ± SEM of Ca^2^+ signals induced by three consecutive applications of 12.5 μM PregS and the effects of four μOR agonists, 0.5 μM DAMGO (red line, n=208), 1 μM Herkinorin (blue, n=173), 0.5 μM TRV130 (brown, n=227) and 1 μM PZM21 (green, n=187). Black trace denotes control experiments with 3 consecutive applications of 12.5 μM PregS, n=108. **(B)** Changes in the relative fluorescence ratio according to the following formula: ΔR_n_/ ΔR_1_ = 100(R_n_ − R_0_)/ (R_1_ − R_0_) normalized to the first pulse of PregS (PS1).

Ca^2+^ signals are a convenient way to monitor the activity of Ca^2+^ permeable ion channels, but they are not linear readouts of channel activity, thus it is difficult to draw quantitative conclusions on the extent of channel inhibition. Therefore we also performed patch clamp experiments to investigate how these biased μOR agonists affect PregS-induced TRPM3 currents in HEK cells (Fig. 3). In these experiments we tested two concentrations for each agonist 100 nM and 1 μM. In each measurement, we applied two pulses of 25 μM PregS; during the first application we also applied the tested agonist and during the second one we applied DAMGO as a control. In all cases application of DAMGO, both 100 nM and 1 μM produced almost full inhibition of both inward (−100 mV) and outward TRPM3 currents (+100 mV). For the three tested substances the inhibitory effects were smaller compared to DAMGO. For 100 nM PZM21, PregS-induced currents were inhibited by 43.4±3.3 % (p=0.0001) at +100 mV and 50.1±8.5 % (p=0.0037) at −100 mV. In the same experiments, 100 nM DAMGO inhibited TRPM3 currents by 79.7±10 % (p<0.00001) for the outward and 72.8±12.9 % (p=0.00011) for the inward currents, n=6 (Fig. 3A). PZM21 at 1 μM induced a more pronounced inhibition 64.9±10.2 % (p<0.00001) for the outward and 62.2±10.8 % (p<0.00001) for the inward currents. In the same experiments 100 nM DAMGO inhibited currents by 93.2±4.5 % (p<0.00001) for the outward and 86.6±1.6 % (p<0.00001) for the inward (−100 mV) currents, n=6 (Fig. 3B). Application of 100 nM Herkinorin had minor effects on PregS-induced currents both at +100 mV, evoking 21.1 ±5 % inhibition (p=0.015) and −100 mV 31±11.2 % 37 % inhibition (p=0.025), while the identical concentration of DAMGO induced 75.2±5.4 % inhibition (p=0.00002) for +100 mV and 71.7±4.4 % (p<0.00001) for −100 mV in the same experiments, n=5 (Fig. 3C). Application of 1 μM Herkinorin had a more pronounced inhibitory effect and lead to current inhibition 53.7±11 % (p=0.00027) for the outward current and 58.8±14 % (p=0.0017) for the inward current. In the same experiments 1 μM DAMGO lead to current inhibition by 89.1±3 % (p<0.00001) for the outward and 83.2±3.9 % (p=0.00007) for the inward currents, n=5 (Fig. 3D). TRV130 behaved similarly; at 100 nM it decreased TRPM3 currents by 45.6±12 % (p=0.0029) for +100 mV and by 48.2±12.8 % (p=0.0019) for -100 mV, compared to 100 nM DAMGO, which inhibited currents by 84.2±6.5 % (p<0.00001) for +100 mV and by 76.9±4.9 % (p<0.00001) for -100 mV, n=6 (Fig. 3E). TRV130 at 1 μM induced a higher level of inhibition, 76.8±7.1 % (p<0.00001) for the outward and 78.9±6 % (p<0.00001) for the inward currents. In the same experiments 100 nM DAMGO inhibited currents on 95.1±2.1 % (p<0.00001) for the outward and 86.7±6.8 % (p<0.00001) for the inward currents, n=6 (Fig. 3F). All biased agonists induced a significantly smaller inhibition than DAMGO in the same experiments (Fig. 3H).

**Fig 3.**
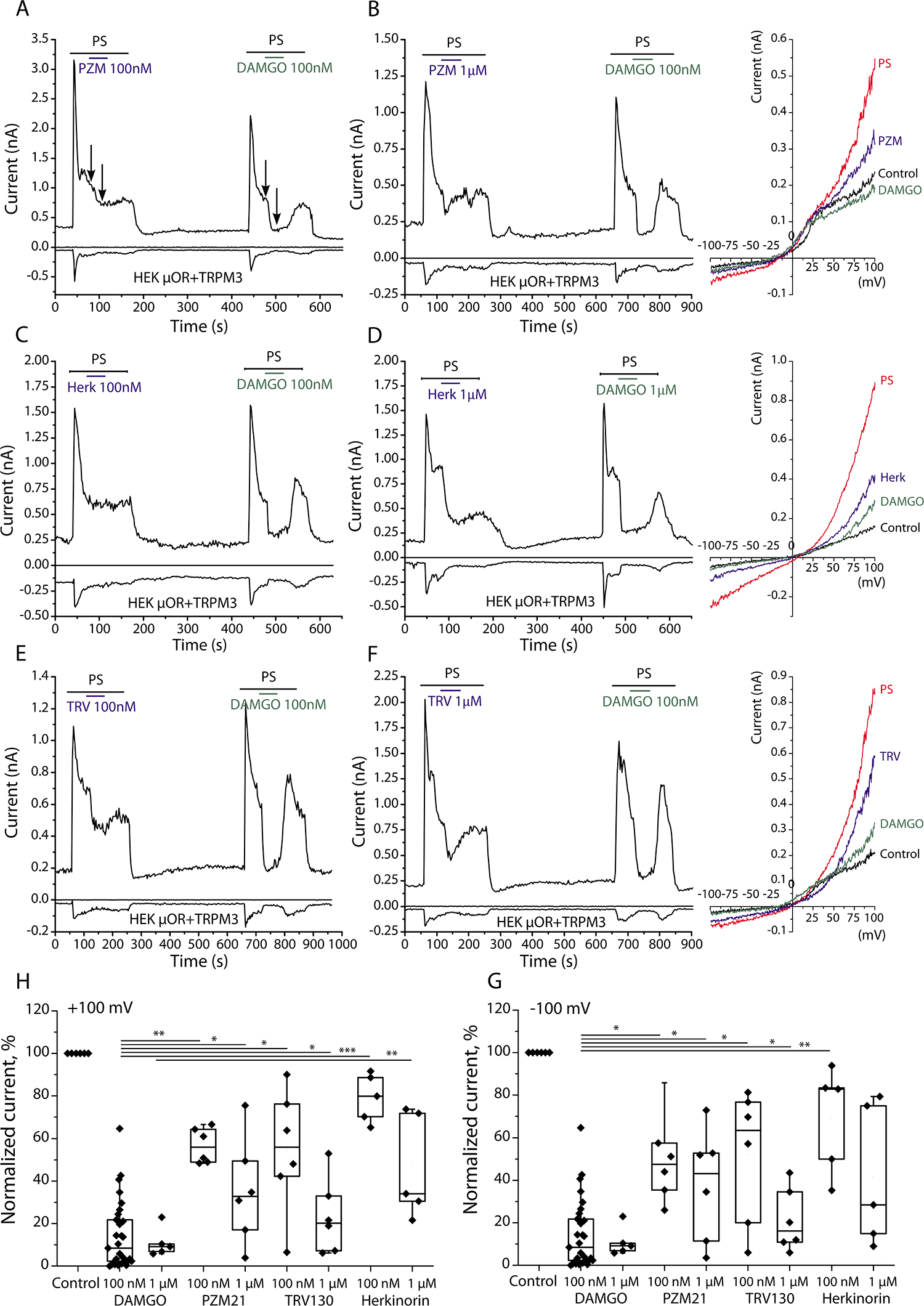
Differential inhibition of PregS-induced TRPM3 currents by μOR agonists in HEK293 cells. Whole-cell patch clamp measurements in HEK293 cells transfected with mTRPM3 channels and μOR were performed as described in Materials and methods. **(A, B)** Representative traces for 100 nM PZM21 (n=6), 1 μM PZM21 (n=6) and 100 nM DAMGO on Preg-S induced current. The applications of 25 μM PregS, DAMGO and PZM21 are indicated by the colored-coded horizontal lines. Right panel shows color-coded I-V curve representative for the corresponding time point measurements. **(C, D)** Representative traces for 100 nM Herkinorin and 1 μM DAMGO (n=5), 1 μM Herkinorin and 1 μM DAMGO (n=5) on PregS-induced current. The applications of 25 μM PregS, DAMGO and Herkinorin are indicated by the colored-coded horizontal lines. Right panel show color-coded I-V curve representative for the corresponding time point measurements. **(E, F)** Representative traces for 100 nM TRV130 (n=6), 1 μM TRV130 (n=6) and 100 nM DAMGO on PregS-induced current. The applications of 25 μM PregS, DAMGO and TRV130 are indicated by the colored-coded horizontal lines. Right panel show color-coded I-V curve representative for the corresponding time point measurements. **(G, H)** Summary for the of DAMGO, PZM21, TRV130 and Herkinorin effects on normalized inward and outward TRPM current densities. Current values were normalized to steady-state PregS (PS) induced currents 2 sec prior to μOR agonists application. Statistical analysis was performed with paired sample t-test *p<0.05, **p<0.01 within each experimental group, data on the graph for 100 nM DAMGO were pooled from different experiments for visualization.

### PZM21 and TRV130 relieve DAMGO-induced inhibition of TRPM3 currents

To test if these new reagents behave as partial agonists we performed patch clamp recordings where TRPM3 was first inhibited by 100 nM DAMGO, then PZM21 or TRV130, was co-applied with DAMGO (Fig 4). We found that 100 nM DAMGO inhibited PregS-induced currents by 97.5±3.2 % for the outward (+100 mV) and 94.9±2.8 % for the inward (−100 mV) currents. When 1 μM PZM21 was co-applied with DAMGO, TRPM3 currents recovered substantially up to 32.7±7.5 % inhibition (p<0.00001 vs. DAMGO inhibition) for the outward (+100 mV) and 33.3 ±8.9 % (p=0.0001 vs. DAMGO inhibition) for the inward (−100 mV) currents, n=9 (Fig. 4A,B). When PZM21 was washed out in the continuous presence of DAMGO, currents returned to close to baseline again (Fig. 4A,B).

**Fig 4.**
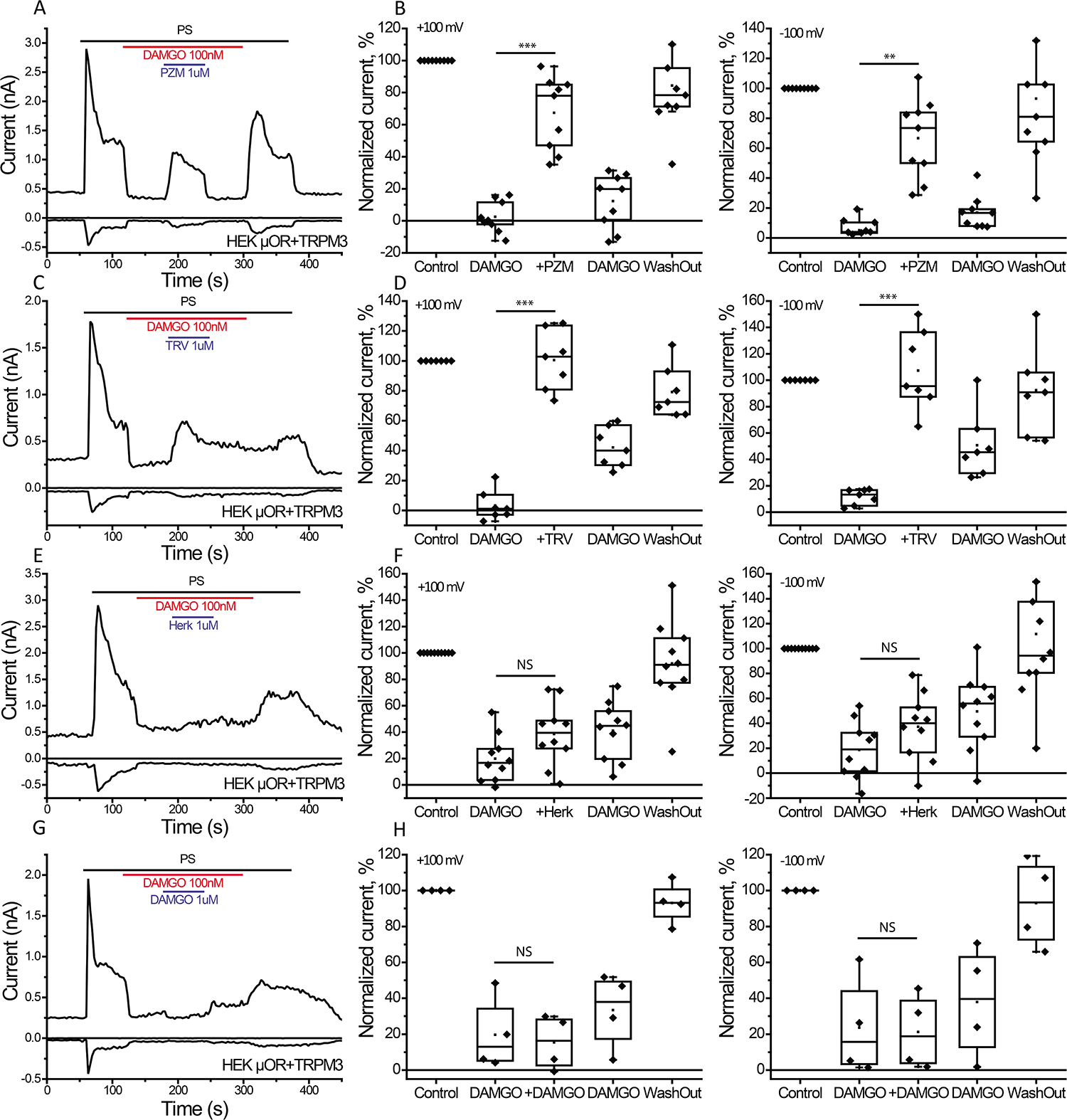
PZM21 and TRV130 oppose the inhibitory effect of DAMGO on PregS-induced TRPM3 currents in HEK293 cells. Whole-cell patch clamp measurements in HEK293 cells transfected with mTRPM3 channels and μOR were performed as described in Materials and methods. **(A)** Representative traces for the effects of PZM21 on PregS current inhibition induced by DAMGO, n=9. The applications of 25 μM Preg-S, 100 nM DAMGO and 1 μM PZM21 are indicated by the colored-coded horizontal lines. **(B)** Summary of normalized current densities for the outward (left) and inward currents (right panel). Inward (−100 mV) and outward currents (+100 mV) were normalized to steady-state Preg-S (PS) induced current 2 sec prior of DAMGO application. Statistical analysis was performed with ANOVA *p<0.05, **p<0.01. **(C)** Representative traces for the effect of TRV130 on PregS current inhibition induced by DAMGO, n=7. The applications of 25 μM PregS (PS) 100 nM DAMGO and 1 μM TRV130 are indicated by the colored-coded horizontal lines. **(D)** Summary of normalized current densities for the outward (left) and inward currents (right panel) for the effect of TRV130. (E) Representative traces for Herkinorin effect on PregS current inhibition induced by DAMGO, n=10. The applications of 25 μM PregS, 100 nM DAMGO and 1 μM Hekinorin are indicated by the colored-coded horizontal lines. **(F)** Summary of normalized current densities for the outward (left) and inward current (right panel) for the effect of DAMGO. **(G)** Representative traces for the effect of 1 μM DAMGO on PregS current inhibition induced by 100 nM DAMGO, n=4. The applications of 25 μM PregS, 100 nM DAMGO and 1 μM DAMGO are indicated by the colored-coded horizontal lines. **(H)** Summary of normalized current densities for the outward (left) and inward current (right panel) for the effect of DAMGO.

Similar effects were observed also for 1 μM TRV130 (Fig. 4C,D). In these experiments DAMGO inhibited PregS-induced currents by 96.8±3.7 % for +100 mV and 88.4±2.2 % for −100 mV, and TRV130 reversed this inhibition to 3.9±7.5 % potentiation (p<0.00001 vs. DAMGO inhibition) at +100 mV and 7.2±11.4 % potentiation (p<0.00001 vs. DAMGO inhibition) at −100 mV, n=7. As opposed to the effect of PZM21, the effect of TRV130 was not quickly reversible (Fig. 4C).

As opposed to PZM21 and TRV130, application of 1 μM herkinorin did not induce a robust recovery from DAMGO-induced inhibition (Fig. 4 E,F). In this set of experiments we observed that 100 nM DAMGO inhibited PregS-induced currents by 81.2±5.6% at +100 mV; after co-application of herkinorin, inhibition decreased to 61.5±7.4%, but this effect was not statistically significant (p=0.63 vs. DAMGO inhibition). At −100 mV DAMGO induced an 81.4±7.2% inhibition, which decreased to 62.8±8.4% after the co-application of herkinorin (p=0.77 vs. DAMGO inhibition, n=10).

As a control we co-applied 1 μM DAMGO in the presence of 100 nM DAMGO which slightly, but not statistically significantly increased the initial inhibitory level (Gig. 4G,H). After the initial 100 nM DAMGO application outward currents were inhibited by 80.4±10% and 1 μM DAMGO it further increased inhibition to 84.6±7.5%. Similarly inward currents changed from 76.4±13.8% to 79.8±10.4% inhibition when 1 μM DAMGO was co-applied with 100 nM DAMGO n=4.

Altogether our data show that PZM21, TRV130 and Herkinorin induced significantly smaller inhibition of TRPM3 currents than the full μOR agonists DAMGO. We also show that PZM21 and TRV130 behave as partial agonists, as they induce recovery from DAMGO-induced inhibition of TRPM3 currents. These results can imply that the β-arrestin pathway is required for full inhibition of TRPM3, or alternatively, that these three G-protein biased compounds are partial agonists on μOR. To differentiate between these alternatives, we tested the effects of activating μOR with these compounds on two well-established G_βγ_ regulated ion channels GIRK2 and N-type Voltage Gated Ca^2+^ channels (Ca_V_2.2).

### PZM21, TRV130 behave as partial μOR agonists in activating GIRK channels

G_βγ_ activation of GIRK channels is one of the classical downstream effects of G_αi_ signaling. We co-expressed Kir3.2 (GIRK2) channels with μOR in HEK cells and in patch clamp experiment we observed that application of the full agonist DAMGO induced strong concentration-dependent activation of GIRK2 channels (Fig 5); 10 nM DAMGO induced average inward current density −16 pA/pF (n=6), 100 nM evoked −47.5 pA/pF (n=18) and 1 μM induced −70.4 pA/pF (n=10). We used 1 μM of the tested reagents that was identical to the maximal tested concentration of DAMGO and for all of them activation of GIRK channels were significantly smaller than that induced by the same concentration of DAMGO; PZM21-induced currents reached −16.3 pA/pF (n=7, p=0.0012 vs. 1 μM DAMGO), for TRV130 −9.3 pA/pF (n=5, p=0.00099 vs. 1 μM DAMGO) and for Herkinorin −25.2 pA/pF (n=5, p=0.041 vs. 1 μM DAMGO).

**Fig 5.**
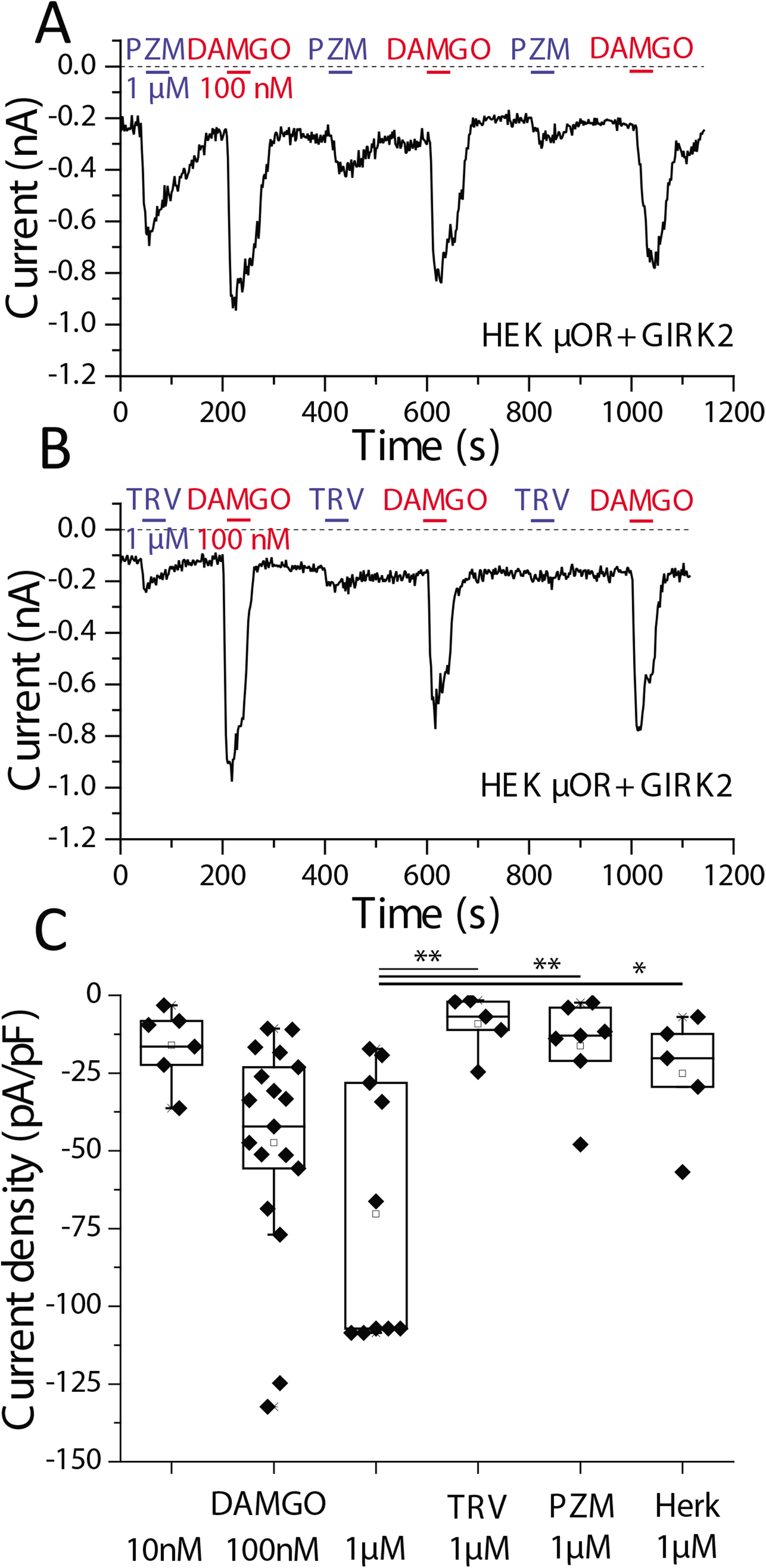
Differential activation of GIRK2 channels by μOR agonists in HEK293 cells. Whole-cell patch clamp measurements in HEK cells transfected with μOR and GIRK2 channels were performed as described in Materials and methods. **(A)** Representative traces for inward (−100 mV) currents for the effects of 1 μM PZM21 and 100 nM DAMGO on GIRK2 currents. **(B)** Representative traces for inward (−100 mV) currents for the effects of 1 μM TRV130 and 100 nM DAMGO on GIRK2 channels. **(C)** Summary of current densities for GIRK2 channels activated by 10 nM DAMGO (n=6), 100 nM DAMGO (n=18), 1 μM DAMGO (n=10) and 1 μM TRV130 (n=5), 1 μM PZM21 (n=7), 1 μM Herkinorin (n=5).

To test the possibility of the partial agonist behavior of PZM21, TRV130 and Herkinorin we co-applied them with DAMGO. After strong DAMGO activation of GIRK2 channels, PZM21 and TRV130 quickly induced significant and prominent reduction of the potassium current amplitude, while Herkinorin did not have a clear inhibitory effect (Fig. 6). In control experiments the application of 1 μM DAMGO slightly potentiated the effect of 100 nM DAMGO (Fig 6). GIRK2 currents induced by 100 nM DAMGO were inhibited upon the application 100 nM PZM21 by 52.6±7.9% (p<0.00001, n=5) and for 1 μM PZM21 GIRK currents were inhibited by 71.4±5.4 % (p<0.00001, n=7). TRV130 (100 nM) inhibited GIRK currents by 51.4±4.6 % (p<0.00001, n=6) and 1 μM TRV130 reduced DAMGO-induced GRIK currents by 73.4±4.8 % (p<0.00001, n=8).

**Fig 6.**
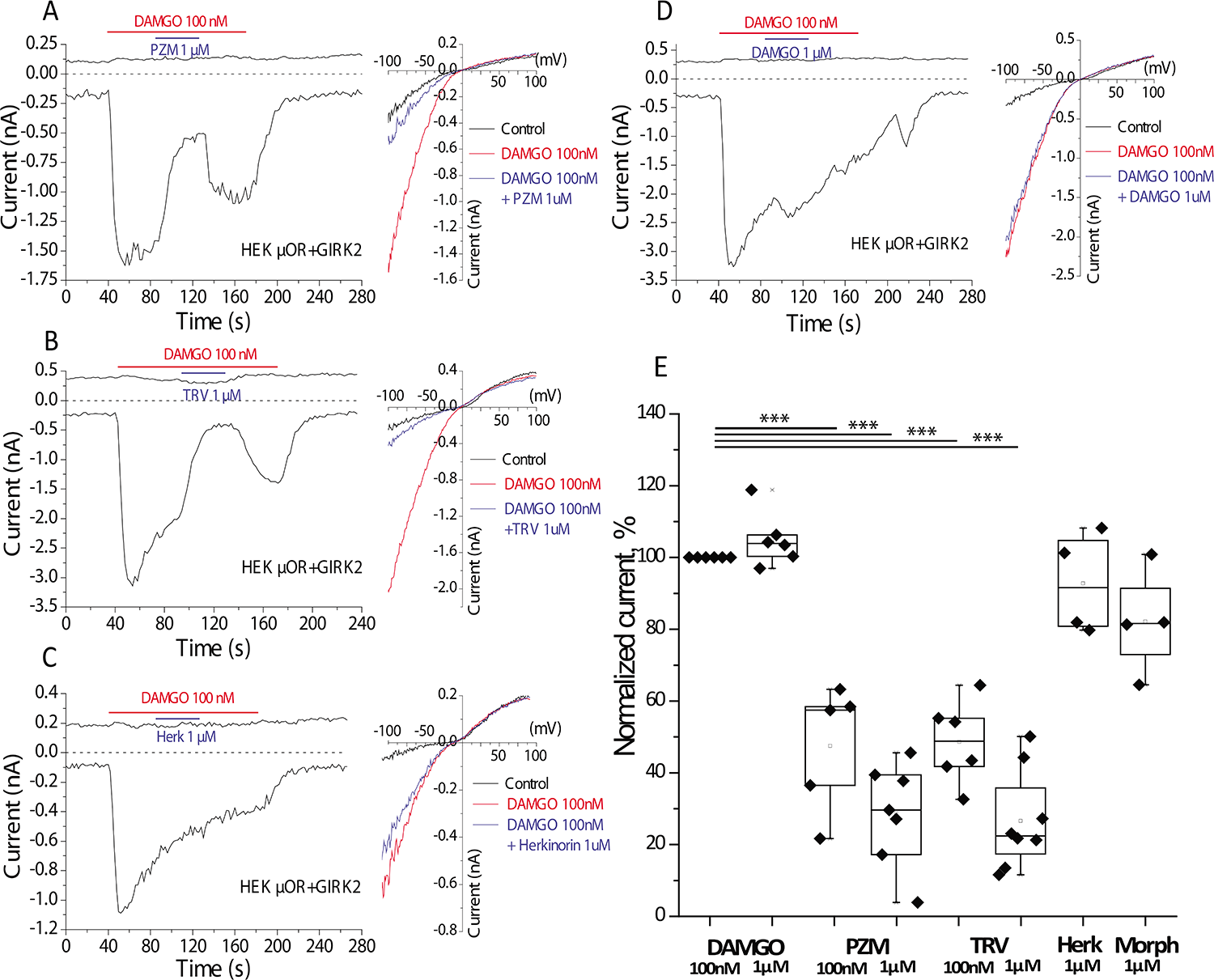
Inhibition of DAMGO-induced GIRK2 currents by PZM21 and TRV130 in HEK293 cells. Whole-cell patch clamp measurements in HEK cells transfected with μOR and GIRK2 channels were performed as described in Materials and methods. **(A)** Representative traces for the effects of 1 μM PZM21 (n=7) on DAMGO-activated GIRK2 currents, n=6. The applications of 100 nM DAMGO and 1 μM PZM21 are indicated by the horizontal lines. Right panel show I-V curve representative for the corresponding time point measurements. **(B)** Representative trace for the effect of 1 μM TRV130 on DAMGO-induced GIRK2 currents, n=8. The applications of 100 nM DAMGO and 1 μM PZM21 are indicated by the horizontal lines. Right panel shows color-coded I-V curves representative for the corresponding time point measurements. **(C)** Representative traces for the effect of 1 μM Herkinorin on DAMGO-induced GIRK2 currents, n=4. The applications of 100 nM DAMGO and 1 μM Herkinorin are indicated by the horizontal lines. Right panel show color-coded I-V curve representative for the corresponding time point measurements. **(D)** Representative traces for the effect of 1 μM DAMGO on GIRK2 currents activated by 100 nM DAMGO, n=6. The applications of 100 nM DAMGO and 1 μM DAMGO are indicated by the colored-coded horizontal lines. Right panel shows color-coded I-V curve representative for the corresponding time point measurements. **(E)** Summary of GIRK2 current densities. Currents induced 100 nM DAMGO were measured before the application, and after wash out of the test compounds; the mean of these two current amplitudes were used as the normalizing point to assess the effects of all tested compounds. Statistical analysis was performed with ANOVA *p<0.05, **p<0.01.

Herkinorin at 1 μM resulted in a current decrease corresponding to 7.2±7.0 % inhibition (p=0.38), n=4, but the currents showed substantial desensitization, therefore this small decrease may not be due to herkinorin (Fig. 6C). To account for this spontaneous current decrease, inhibition levels were calculated by taking the average of GIRK currents before and after application of the test compounds as the control level for all tested compounds. In a control experiment application of 1 μM DAMGO on top of 100 nM DAMGO lead to a slight potentiation 5.0±3.1 % (p=0.49), n=6 (Fig. 6D). Morphine (1 μM) did not induce a significant inhibition of DAMGO-induced GIRK currents (Fig. 6E).

### G-protein biased μOR agonists inhibit N-type voltage gated channels less efficiently than the full agonist DAMGO

N-type voltage gated Ca^2+^ channels (Ca_v_2.2) are also classical targets for G_βγ_ subunits. In cells co-transfected with μOR and Ca_v_2.2 channel subunits we also recorded strong inhibition of the Ba^2+^ current by 1 μM DAMGO 34.4±4.7 %, p<0.00001, n=12) but for the two other tested reagent’s level of inhibition was less prominent, 1 μM PZM21 demonstrated inhibition 17.5±8 % (p=0.061, n=5) and 1 μM TRV130 13.8±2.3 % (p=0.009, n=8), Fig 7. Both the effects of PZM21 (Fig. 7C) and TRV130 (Fig. 7F) were significantly smaller than that of DAMGO.

**Fig 7.**
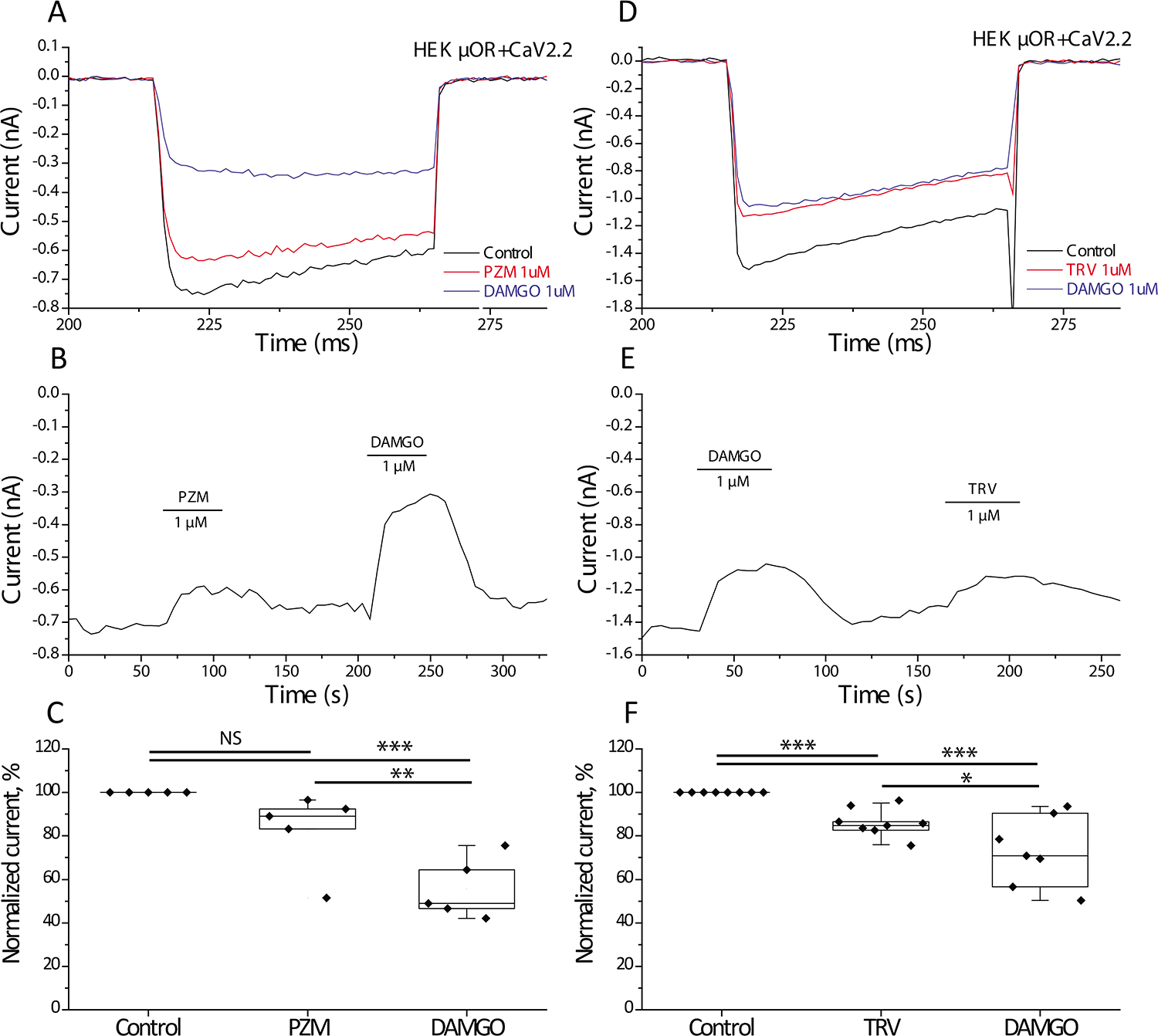
The effects of pOR agonists on Ca_V_2.2 channels in HEK293 cells. Whole-cell patch clamp measurements in HEK293 cells transfected with μOR and Ca_V_2.2 channel subunits were performed as described in Materials and methods. **(A)** Representative traces of Ca_V_2.2 barium currents activated by voltage steps from *V*_h_ −80 mV to +10 mV and the inhibitory effect of 1 μM PZM21 and 1 μM DAMGO. **(B)** Time course of the I_Ba_ in the presence of PZM21 and DAMGO. **(C)** Summary of normalized currents for 1 μM PZM21 and 1 μM DAMGO (n=5). **(D)** Representative traces of Ca_V_2.2 barium currents activated by voltage steps from *V*_h_ −80 mV to +10 mV and the inhibitory effect of 1 μM TRV130 and 1 μM DAMGO. **(E)** Time course of the I_Ba_ in the presence of TRV130 and DAMGO. **(F)** Summary of normalized current for 1 μM TRV130 and 1 μM DAMGO (n=8).

Our data show that G-protein biased μOR agonists were less efficient than the full agonist DAMGO in modulating three different ion channel targets to G_αi_-G_βγ_ signaling. We next tested if the G-protein biased agonists were also less efficient than DAMGO on G-protein dissociation, using a FRET-based assay.

### G-protein μOR biased agonists are less efficient than DAMGO in inducing G-protein dissociation

To assess the dissociation level of G_α_ and G_βγ_ during μOR activation we used a FRET-based assay in cells transfected an mTurqoise2-tagged G_αi1_ (FRET donor), a Venus-tagged G_γ2_ (FRET acceptor) and G_β1_ expressed from a tricistronic vector (van Unen et al., 2016), and with μOR. The reduction of FRET signal in these experiments indicates dissociation of the G_α_-G_βγ_ heterotrimer complex, or a change in relative conformation between the FRET donor and acceptor. In this experiment cells treated with different DAMGO concentrations demonstrated clear antiparallel signals, an increase at 480 nm and a decrease at 530 nm emission wavelengths indicating a decrease in FRET (Fig 8A,C). For 10 nM DAMGO, the FRET signal showed a slight 0.48±0.12 % decrease (p=0.0068, n=4), 100 nM evoked 2.49±0.36 % (p<0.00001, n=8), and 1 μM DAMGO resulted in 3.92 ±0.27 % (p<0.00001, n=36) decrease in the FRET signal. We observed significantly weaker agonist responses in FRET experiments for 1 μM PZM21 (1.58±0.41 % decrease, p=0.0017, n=8), 1 μM TRV130 (1.03±0.17 % decrease, p=0.00013, n=6) and 1 μM herkinorin (1.52±0.32 % decrease, p<0.00001, n=5). These data indicate a smaller level of dissociation of G_αi1_ and G_βγ_ compared to the effect of the full μOR agonist DAMGO by these agonists. As 1 μM herkinorin did not evoke a clear reversal of DAMGO-induced inhibition of TRPM3 and DAMGO-induced activation of GIRK, we also tested higher concentrations of this drug in the FRET assay. For 10 μM herkinorin we observed 0.88±0.14 % of FRET signal decrease (p<0.0018, n=8) an effect very similar to that induced by 1 μM herkinorin. The effects of PZM21, TRV130 and herkinorin were significantly smaller than that induced by 1 μM DAMGO (Fig. 8C).

**Fig 8.**
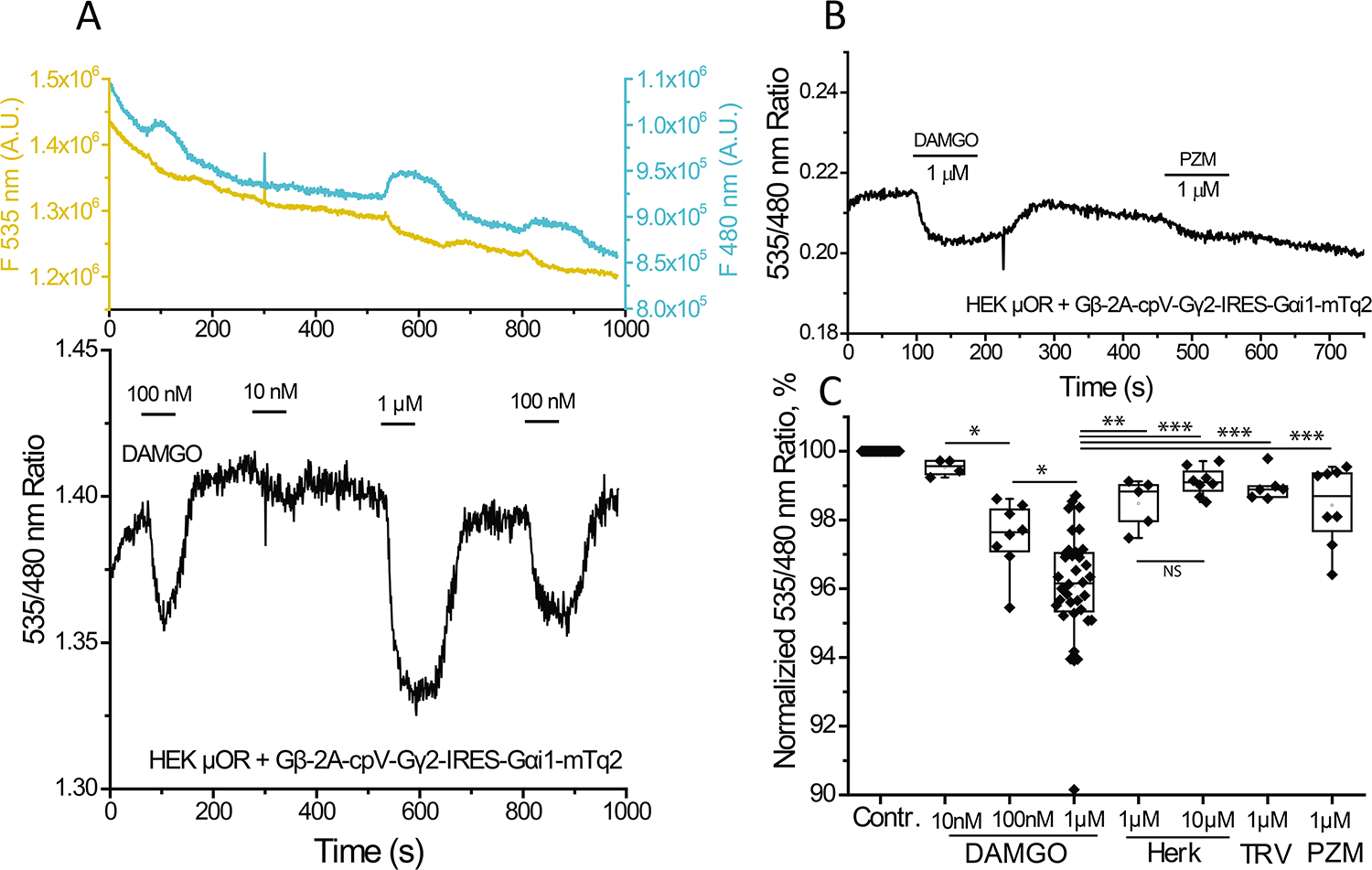
Dissociation level of Gα and Gβ_γ_ evoked by μOR agonists in FRET experiment. Fluorescence measurements were performed in HEK293 cells transfected with mTurqoise2-tagged G_αi1_ (FRET donor), a Venus-tagged G_γ2_ (FRET acceptor) and G_β1_ expressed from a tricistronic vector (Gβ-2A-cpV-G_γ_2-IRES-Gαi1-mTq2) and μOR as described in the Methods section. **(A)** Upper panel, fluorescence traces detected at 480 for FRET donor (blue line) and 535 nm for FRET acceptor (yellow line); excitation wavelength was 430 nm. Lower panel shows ratio of the individual traces (FRET signal). The applications of 10 nM, 100 nM and 1 μM DAMGO are indicated by the horizontal lines. **(B)** Similar measurement on cells tested with 1 μM DAMGO and 1 μM PZM21. **(D)** Normalized responses of FRET signals evoked by the applications of 10 nM DAMGO (n=4), 100 nM DAMGO (n=8) and 1 μM DAMGO (n=36), 1 μM TRV130 (n=6), 1 μM PZM21 (n=8), 1 μM Herkinorin (n=5) and 10 μM Herkinorin (n=8). All fluorescence ratio traces were normalized to the ratio level before the application of the stimulus.

In conclusion we demonstrate that the recently described G_αi_-biased agonists PZM21 and TRV130 behave as weak partial agonists using three ion channel targets for the G_βγ_ subunits as a functional readout.

## DISCUSSION

Most available opioid medications activate the G_αi_-coupled μOR-s; ligand binding to these receptors leads to the dissociation of G_α_ and G_βγ_ and also induces β-arrestin recruitment. Both G-protein subunits have their own effectors; G_αi_ inhibits adenylate cyclase (AC), and thus decrease cAMP level. G_βγ_ subunits have multiple targets including G protein-coupled inwardly-rectifying potassium channels (Kir3) (Logothetis et al., 1987) and voltage-gated Ca^2+^ channels (N-type and P/Q-type) (Currie, 2010). A recently described target of G_i_-signaling is the TRPM3 ion channel, which is robustly inhibited by G_βγ_ subunits (Badheka et al., 2017; Dembla et al., 2017; Quallo et al., 2017). Non-ion channel targets of G_βγ_ include phosphoinositide 3-kinase-γ (PI3Kγ) and mitogen-activated protein kinases (Khan et al., 2013), (Clapham and Neer, 1997).β-arrestin 1 (arrestin 2) and β-arrestin 2 (arrestin 3) are important regulators of G-protein signaling. They were originally described as mediators of agonist-induced desensitization and internalization (DeWire et al., 2007), and they were presumed to be responsible for many side effects of opioids, such as tolerance and respiratory depression. Recently β-arrestins have emerged as independent signaling mediators regulating various effectors (DeWire et al., 2007), including ion channels in the TRP family (Liu et al., 2017). We initiated this study to determine the possible role of the β-arrestin pathway in regulating TRPM3 channels, the newly described targets of G_βγ_ signaling.

We found that all three G-protein biased μOR agonists, PZM21, TRV130 and Herkinorin induced significantly smaller inhibition of TRPM3 channels compared to the full agonist DAMGO both in Ca^2+^ imaging and patch-clamp experiments. PZM21 and TRV130 also reversed the inhibitory effect of DAMGO when co-applied with this full agonist. These results may indicate the involvement of the β-arrestin pathway in TRPM3 regulation. Initially designed as control experiments, we also found that all these reagents induced a weaker GIRK channel activation than DAMGO. Similar to TRPM3 channels, PZM21 and TRV130, but not herkinorin robustly opposed the effect on DAMGO on GIRK channels when co-applied with this full agonist. Voltage gated Ca_V_2.2 channels were also inhibited less by PZM21 and TRV130 compared to DAMGO. These results could indicate two possibilities; either the β-arrestin pathway is required for the activation of all three ion-channel targets of Gβ_γ_, or PZM21, TRV130 and herkinorin act as partial agonists in G-protein signaling. To differentiate between these two possibilities, we reexamined the effects of these compounds on G-protein dissociation. We performed FRET experiments and found that all tested reagents induced weaker G_α_-G_βγ_ dissociation than DAMGO, indicating that PZM21 and TRV130 behave as weak partial agonists when regulating ion channel targets for G_βγ_ subunits.

All three compounds tested here were developed as G-protein biased μOR agonist to avoid recruitment of β-arrestin with the promise of novel pain medications with fewer side effects. Herkinorin demonstrated strong antinociceptive effects, similar to that of morphine, in several behavioral assays without inducing tolerance (Lamb et al., 2012). At the cellular level it was demonstrated that herkinorin did not recruit β-arrestin 2 and did not promote internalization, but had G_α_ agonist activity similar, or stronger than that of DAMGO, detected by a MAP kinase phosphorylation assay (Groer et al., 2007).

TRV130 was also shown to be effective in a variety of behavioral tests with efficacy and potency similar to that of morphine, while inducing less constipation and respiratory suppression compared to similar doses of morphine (DeWire et al., 2013). As a biased agonist it strongly activated G_αi_ (cAMP assay) without recruitment of β-arrestin 2 (enzyme complementation assay) (DeWire et al., 2013). These findings were confirmed by another independent study on TRV130 (Mori et al., 2017).

The newest G_αi_-biased μOR agonist PZM21 was identified based on computational docking of over 3 million compounds to the structure of μOR, and subsequent structure based optimization (Manglik et al., 2016). PZM21 is structurally unrelated to known opioids; it was shown to induce G-protein activation using various high throughput optical assays, but it induced no recruitment of β-arrestin 2, with very low level of receptor internalization. In behavioral experiments, PZM21 had antinociceptive properties in multiple animal tests with significantly less adverse side effects than known opioids (Manglik et al., 2016). Manglik et al also compared herkinorin and TRV130 to the effects of PZM21, DAMGO as well as morphine is several high throughput cellular assays. They concluded that PZM21 is a G-protein biased, potent, selective, and efficacious μOR agonist; TRV130 behaved similar to PZM21, but they found that herkinorin induced substantial β-arrestin recruitment.

What is the explanation of the difference between our data and those of Manglik et al? One possible explanation is that the different assays used to assess G-protein activity have different sensitivities and dynamic ranges. The high throughput optical assays, such as glosensor are not linear readouts of cAMP levels, and while the assay is highly sensitive at low levels of activity, it is possible that at high activities such that induced by DAMGO it may saturate, which could overestimate the maximal effect of a weaker agonist. Indeed while PZM21 was designated as efficacious in G-protein stimulation, it induced between 70 % and ~100 % of the effect of DAMGO depending on the assay (Manglik et al., 2016). Ionic currents measured in patch clamp experiments on the other hand, are linear readouts of channel activity, which do not saturate, thus this technique is more sensitive at high activity levels.

In our FRET-based G-protein dissociation experiments all three agonists induced well below 50% of the signal induced by DAMGO. In our ion channel experiments, the difference between DAMGO and the biased agonists were variable; depending on the channel, and the concentrations used, but it was again, generally well below the effect induced by DAMGO. We used 100 nM and 1 μM for all compounds; 1 μM was a saturating concentration for all agonists, whereas 100 nM induced maximal or close to maximal responses in most assays (Manglik et al., 2016). Our most definitive data showing that these compounds act as partial agonists is the clear reversal of the effect of DAMGO on TRPM3 and GIRK channels by PZM21 and TRV130 (Figs. 4 and 6). Herkinorin at 100 nM or 1 μM did not induce a robust reversal of the effect of DAMGO on TRPM3 currents (Fig. 4) and on GIRK channels (Fig. 6), but the marked desensitization of both currents makes a small effect difficult to assess. Because of this ambiguity, we also tested the effect of 10 μM herkinorin in our FRET based assay, and its effect was indistinguishable from that induced by 1 μM herkinorin. It is not clear why herkinorin behaves differently from PZM21 and TRV130 in reversing the effects of DAMGO, but we have not further investigated this difference.

Our observations are in line with a recent publication showing that PZM21 acts as a partial agonist with substantially lower efficacy than DAMGO, using a BRET-based assay to detect the dissociation of G_αi1_ and G_γ2_ (Hill et al., 2018). Using the same assay they also found that morphine had a higher efficacy than PZM21, but lower than that of DAMGO (Hill et al., 2018). Our data are also compatible with another recent publication using a conformational biosensor binding to the purified μOR showing that PZM21 acted as a partial agonist, with lower efficacy than morphine, or phentanyl, two clinically used opiods (Livingston et al., 2018).

In conclusion, our data indicate that the G-protein biased PZM21 and TRV130 act as weak partial μOR agonists, when signaling to ion channel targets.

## METHODS

### DRG neuron isolation and culture

Animal procedures were approved by the Institutional Animal Care and Use Committee at Rutgers New Jersey Medical School. DRG neurons were isolated from adult mice of either sex (2–4 months old) using the slightly modified protocol based on (Malin et al., 2007), as described previously (Lukacs et al., 2007; Yudin et al., 2016). Briefly, anesthetized animals were perfused via the left ventricle with ice-cold Hank’s buffered salt solution (HBSS; Invitrogen) followed by decapitation. DRGs were collected from all spinal segments after laminectomy and maintained in ice-cold HBSS during the isolation. After isolation and trimming of dorsal and ventral roots, ganglia were incubated in an HBSS-based enzyme solution containing 2 mg/ml type I collagenase (Worthington) and 5 mg/ml Dispase (Sigma) at 37°C for 25–30 min, followed by repetitive trituration for dissociation. After centrifugation at 80 × g for 10 min, cells were resuspended and plated on round coverslips pre-coated with poly-l-lysine (Invitrogen) and laminin (Sigma), allowed to adhere for 1 hr and maintained in culture in DMEM/F12 supplemented with 10% FBS (Thermo Scientific), 100 IU/ml penicillin and 100 μg/ml streptomycin and were kept in a humidity-controlled tissue-culture incubator with 5% CO_2_ at 37°C for 12–36 hr. before experiments.

### HEK293 cells maintenance

Human Embryonic Kidney 293 (HEK293) cells were purchased from American Type Culture Collection (ATCC), Manassas, VA, (catalogue # CRL-1573), RRID:<u>CVCL_0045</u>. Passage number of the cells was monitored, and cells were used up to passage number 25–30, when a new batch of cells was thawed with low passage number. The cells were maintained in minimal essential medium (MEM) (Life Technologies, Carlsbad, CA, USA) supplemented with 10% (v/v) fetal bovine serum (FBS), 100 IU/ml penicillin and 100 μg/ml streptomycin in an incubator (37°C in 5% CO_2_).

### Ca^2+^ imaging

Ca^2+^ imaging measurements were performed with an Olympus IX-51 inverted microscope equipped with a DeltaRAM excitation light source (Photon Technology International, Horiba), as described earlier (Lukacs et al., 2013). Briefly, DRG neurons or HEK cells were loaded with 1 μM fura-2 AM (Invitrogen) at 37°C for 40-50 min before the measurement, and dual-excitation images at 340 and 380 nm excitation wavelengths were detected at 510 nm with a Roper Cool-Snap digital CCD camera. Measurements were conducted in the same bath solution we used for whole-cell patch clamp recording of TRPM3 channels. Raw data analysis was performed using the Image Master software (PTI). We calculated the normalized change in the relative 340/380 ratio, according to the following formula: ΔR_n_/ ΔR_1_ = 100(R_n_ − R_0_)/ (R_1_ − R_0_), where R_n_ is the maximum amplitude during application of agonist (n=1 for the first PregS pulse and n=2 for the second PregS pulse) and R_0_ is the basal level of fluorescence ratio before stimulation.

### TRPM3 channel electrophysiology

Whole-cell patch clamp measurements were performed as described earlier (Badheka, 2015). The cells were transiently transfected with cDNA encoding the mouse TRPMα2 (mTRPM3α2) splice variant of TRPM3, in the bicistronic pCAGGS/IRES-GFP vector (Oberwinkler et al., 2005; Vriens et al., 2011), and μOR (*cDNA Resource Center, catalogue* #OPRM10FN0) in ratio 1:1 using the Effectene reagent (Qiagen) according manufacturer’s protocol and used in experiments 48-72 hours later. Measurements were carried out on GFP positive cells, in an extracellular solution containing (in mM) 137 NaCl, 5 KCl, 1 MgCl_2_, 2 CaCl_2_, 10 HEPES and 10 glucose, pH 7.4. The intracellular solution contained (in mM) 140 potassium gluconate, 5 EGTA, 1 MgCl_2_, 10 HEPES, and 2 NaATP, pH 7.3. Patch clamp pipettes were prepared from borosilicate glass capillaries (Sutter Instruments) using a P-97 pipette puller (Sutter Instrument) and had a resistance of 4-6 MΩ. In all experiments after formation of gigaohm-resistance seals, the whole-cell configuration was established and currents were recorded using a ramp protocol from −100 mV to +100 mV over 500 ms preceded by a −100 mV step for 200 ms; the holding potential was −60 mV, and this protocol was applied once every 2 seconds. The currents were measured with an Axopatch 200B amplifier, filtered at 5 kHz, and digitized through Digidata 1440A interface. In all experiments, cells that had a passive leak current more than 100 pA were discarded. Data were collected and analyzed with the PClamp10.6 (Clampex) acquisition software (Molecular Devices, Sunnyvale, CA), and further analyzed and plotted with Origin 8.0 (Microcal Software Inc., Northampton, MA, USA).

### GIRK2 channel (Kir3.2) electrophysiology

All recordings were performed in HEK293 cells after transient transfection with plasmids Kir3.2 (gift of Dr. Tooraj Mirshahi), μOR and pEYFP using the Effectene reagent (Qiagen) according manufacturer’s protocol with the ratio of cDNA of 1:1:0.1. Patch-clamp recordings were performed 48–72 hours after transfection. Measurements were carried out on YFP positive cells, in an extracellular solution containing (in mM) 137 KCl, 5 NaCl, 1 MgCl_2_, 2 CaCl_2_, 10 HEPES and 10 glucose, pH 7.4. Electrodes (resistance, 4-6 megaohms) were filled with pipette solution containing (in mM) 130 KCl, 2 MgCl_2_, 10 HEPES, 5 Na-EGTA, 3 NaATP, and 0.5 NaGTP (pH 7.25). After cells were voltage-clamped in the conventional whole-cell configuration the currents were recorded using a protocol starting with a step to −100 mV for 50 ms then ramp up to +100 mV over 200 ms and was applied every 2 seconds; the holding potential was −40 mV. The currents were measured with an Axopatch 200B amplifier, filtered at 5 kHz, and digitized through Digidata 1440A interface.

### Ca_V_2.2 electrophysiology

All plasmids with Ca_V_2.2 channels and auxiliary subunits in pMT2 vector were purchased from Addgene (Raghib et al., 2001). All recording were performed in HEK293 cells after transient transfection with plasmids Ca_V_2.2-GFP (*catalogue* #58737), β_1b_-mCherry (*catalogue* # 89892), α_2_δ_1_ subunits (*catalogue* # 58726) and μOR using the Effectene reagent (Qiagen) according manufacturer’s protocol with the ratio of calcium channel subunits and μOR in a ratio of 1:1:1:1. Patch-clamp recordings were performed 48-72 hours after transfection. Transfected cells were visually identified using fluorescence of GFP and mCherry. The whole cell Ca^2+^ currents, carried by barium (*I*_Ba_), were recorded using an extracellular recording solution consisting of (in mM) 30 tetraethylammonium chloride, 100 NaCl, 5 CsCl, 1 MgCl_2_, 10 BaCl_2_, 10 glucose, and 10 HEPES (pH 7.4). Electrodes (resistance, 4-6 megaohms) were filled with pipette solution containing (in mM) 120 CsCl, 2 MgCl_2_, 10 HEPES, 10 EGTA, 4 NaATP, and 0.35 NaGTP (pH 7.25). After cells were voltage-clamped in the conventional whole-cell configuration the currents were recorded using a voltage step protocol from −80 to +10 mV (50 ms) which was applied once every second. The currents were measured with an Axopatch 200B amplifier, filtered at 5 kHz, and digitized through a Digidata 1440A interface. Series resistance was partially compensated using the Axopatch circuitry (about 75–80% prediction and correction; 10 μs lag). Offline leak and capacitive current subtraction used P/4 protocol with the leak pulses applied following the test pulses was carried out using the Clampfit 10.6 software (Molecular Devices).

### FRET-based monitoring of G_αi1_ activation

FRET measurements were performed as described earlier (Borbiro et al., 2015). Briefly, HEK cells were co-transfected with the Gβ-2A-cpV-Gγ2-IRES-Gαi1-mTq2 (Addgene, *catalogue* #69623) (van Unen et al., 2016) and μOR. Fluorescence was detected using a photomultiplier-based dual-emission system mounted on an inverted Olympus IX-71 microscope equipped with 40X oil objective (numerical aperture 1.3). Excitation light (430 nm) was provided by a DeltaRAM light source (Photon Technology International, PTI). Emission was measured at 480 and 535 nm using two interference filters and a dichroic mirror to separate the two wavelengths. Raw data were analyzed with the Felix3.2 program (PTI).

### Application of solutions and reagents

Coverslips with attached cells were placed in a recording bath (volume ~200 μL) which was continually perfused with fresh extracellular solution at a flow rate of ~3–4 ml/min from gravity-fed reservoirs, and viewed using an Olympus IX-51 inverted microscope. Pregeneolone sulfate (PregS) was purchased from Sigma, DAMGO and Herkinorin were purchased from Abcam, PZM21 from MedChem Express, and TRV130 from Adooq Bioscience. DAMGO and morphine stock were dissolved in H_2_O. PregS, Herkinorin, PZM21 and TRV130 were dissolved in DMSO. All stock dilutions were prepared on the day of the experiment.

### Statistics

Data analysis was performed in Excel and Microcal Origin. The normality of the data was verified with the Kolmogorov-Smirnov test. Data were analyzed with t-test, or Analysis of variance with Bonferroni *post-hoc* test *p<0.05, **p<0.01, ***p<0.005. Data are displayed as box-whisker and scatter plots for most figures; mean values are described in the results section for most measurements.

## Acknowledgements

We thank Drs. Veit Flockerzi and Stephan Phillips for the mouse TRPM3a2 clone; the Kir3.2 clone was a kind gift from Dr. Tooraj Mirshahi. This work was supported from NIH grants NS 055159 and GM 093290.

